# Dynamic distortion of orientation representation after learning in the mouse primary visual cortex

**DOI:** 10.1101/2021.03.25.437032

**Authors:** Julien Corbo, John P. McClure, O. Batuhan Erkat, Pierre-Olivier Polack

**Affiliations:** Center for Molecular and Behavioral Neuroscience. Rutgers University – Newark. 197 University Avenue. Newark, NJ 07102. USA; Behavioral and Neural Sciences Graduate Program, Rutgers University – Newark. 197 University Avenue. Newark, NJ 07102. USA

## Abstract

Learning is an essential cognitive mechanism allowing behavioral adaptation through adjustments in neuronal processing. It is associated with changes in the activity of sensory cortical neurons evoked by task-relevant stimuli. However, the exact nature of those modifications and the computational advantages they may confer are still debated. Here, we investigated how learning an orientation discrimination task alters the neuronal representations of the cues orientations in the primary visual cortex (V1) of male and female mice. When comparing the activity evoked by the task stimuli in naïve mice and mice performing the task, we found that the representations of the orientation of the rewarded and non-rewarded cues were more accurate and stable in trained mice. This better cue representation in trained mice was associated with a distortion of the orientation representation space such that stimuli flanking the task-relevant orientations were represented as the task stimuli themselves, suggesting that those stimuli were generalized as the task cues. This distortion was context dependent as it was absent in trained mice passively viewing the task cues and enhanced in the behavioral sessions where mice performed best. Those modifications of the V1 population orientation representation in performing mice were supported by a suppression of the activity of neurons tuned for orientations neighboring the orientations of the task cues. Thus, visual processing in V1 is dynamically adapted to enhance the reliability of the representation of the learned cues and favor generalization in the task-relevant computational space.

**Significance statement:** Performance improvement in a task often requires facilitating the extraction of the information necessary to its execution. Here, we demonstrate the existence of a suppression mechanism that improves the representation of the orientations of the task stimuli in the V1 of mice performing an orientation discrimination task. We also show that this mechanism distorts the V1 orientation representation space, leading stimuli flanking the task stimuli orientations to be generalized as the task stimuli themselves.

## Introduction

Animals benefit from optimizing their performance at tasks they execute regularly. Learning not only induces changes in the neuronal activity of brain regions involved in the behavioral engagement and motor skills improvement (16), it also affects early sensory processing (26, 67). In the mouse primary visual cortex (V1), learning a detection or a discrimination task involving oriented stimuli is linked to several adaptations of early visual processing. At the neuronal level, positive changes were found in the amplitude of the responses (29, 30), the number of responsive neurons (30, 53), the selectivity of individual neurons (28–30, 36, 53) as well as a reduction of the responses’ trial-to-trial variability (30, 36). At the population level, learning increases the discriminability of the neural responses to the task stimuli (36, 53) and can solidify the reliability of ensembles co-activated by those cues (11). Most studies so far have focused on semantic representations of the stimuli during the task, independently of their features (e.g. Go and NoGo cues; (36, 53)), or on changes mapped onto a feature representation space based on discretized, category-like neurons orientation preferences (28, 30, 36). Although some studies suggested more finely-resolved changes for neurons preferring specific bands of orientations (28, 56), little is known about the impact of learning on the representation of the orientations of the task-relevant and non-relevant stimuli by the population of V1 neurons. Yet, encoding the orientation of visual stimuli is central to the computational role of V1. Indeed, orientation (and direction) coding is a well-established feature of V1 neurons (32), stable through time (17, 43) and across the variations of other features like contrast (3, 6), temporal frequency (47), and spatial frequency (34). Orientation coding is embedded in the network connectivity, as the tuning of a neuron results from the specific inputs it receives (13, 35, 39, 64). Hence, the tuning properties of neurons can be used to access feature-related information in a population. Indeed, a decoder for orientation (blind to the tuning of the neurons) will naturally weigh the contribution of neurons in a way that reflects their preference (6). Inversely, using the tuning properties of cells can provide for an efficient population code (58, 70). To investigate the population representation of visual stimuli in the orientation space, we performed calcium imaging in the layer 2/3 of the mouse V1 before and after training in a simple Go/NoGo discrimination task and mapped the evoked neuronal activity as a function of the neurons’ preferred orientations. We found that the V1 representations of the task’s orientations were more precise and reliable in trained mice than in naïve mice. This sharper representation of the task cues was associated with a distortion of the orientation space that led stimuli having orientations flanking the task cues to be represented as the task stimuli themselves. The modifications of the representation of the visual stimuli in the orientation space were context dependent as they were absent in trained mice disengaged from the task.

## Material and methods

All the procedures described below have been approved by the Institutional Animal Care and Use Committee (IACUC) of Rutgers University-Newark, in agreement with the Guide for the Care and Use of Laboratory Animals (National Research Council, 2011).

### Surgery

#### Head-bar implants

10 minutes after systemic injection of an analgesic (carprofen, 5 mg per kg of body weight), adult (3-6 months old) male and female Gad2-IRES-Cre (Jackson stock #019022) x Ai9 (Jackson stock #007909) mice were anesthetized with isoflurane (5% induction, 1.5% maintenance) and placed in a stereotaxic frame. Body temperature was kept at 37°C using a feedback-controlled heating pad. Pressure points and incision sites were injected with lidocaine (2%). Eyes were protected from desiccation with artificial tear ointment (Dechra). Next, the skin covering the skull was incised and a custom-made lightweight metal head-bar was glued to the skull using Vetbond (3M). In addition, a large recording chamber capable of retaining the water necessary for using a water-immersion objective was built using dental cement (Ortho-Jet, Lang). Mice recovered from surgery for 5 days, during which amoxicillin was administered in drinking water (0.25 mg/mL).

#### AAV virus injection

After recovery from the head-bar surgery, mice were anesthetized using isoflurane as described above. A circular craniotomy (diameter = 3 mm) was performed above V1. The AAV vector AAV1.eSyn.GCaMP6f.WPRE.SV40 (UPenn Vector Core) carrying the gene of the fluorescent calcium sensor GCaMP6f was injected at three sites 500 μm apart around the center of V1 (stereotaxic coordinates: −4.0 mm AP, +2.2 mm ML from bregma) using a MicroSyringe Pump Controller Micro 4 (World Precision Instruments, WPI) at a rate of 30 nl/min. Injections started at a depth of 550μm below the pial surface and the tip of the pipette was raised in steps of 100μm during the injection, up to a depth of 200μm below the pial surface. The total volume injected across all depths was 0.7 μl. After removal of the injection pipette, a 3-mm-diameter coverslip was placed over the dura, such that the coverslip fits entirely in the craniotomy and was flush with the skull surface. The coverslip was kept in place using Vetbond and dental cement. Mice were left to recover from the surgery for at least 3 weeks to obtain a satisfactory gene expression.

### Imaging

#### Calcium imaging setup

During the last week of recovery, mice were trained to stay on a spherical treadmill consisting of a ball floating on a small cushion of air that allowed for full 2D movement (52). During three daily twenty-minute sessions, the mouse head-bar was fixed to a post holding the mouse on the apex of the spherical treadmill. Ball motion was tracked by an IR camera taking pictures of the ball at 30 Hz. Eye motion was monitored at 15 Hz using a second IR camera imaging the reflection of the eye on an infrared dichroic mirror. Functional imaging was performed at 15 frames per second using a resonant scanning two-photon microscope (Neurolabware) powered by a Ti-Sapphire Ultra-2 laser (Coherent) set at 910 nm. The microscope scanning mirrors were hermetically sealed in a chamber to bring the scanning hum below the room ambient noise (< 59 dBA). The laser beam was focused 200 microns below the cortical surface using a 16×, 0.8 NA Nikon water-immersion objective. The objective was tilted 30 degrees such that the objective lens was parallel to the dura surface. Laser power was kept below 70 mW. Frames (512×796 pixels) were acquired using the software Scanbox developed by Neurolabware.

#### Naïve imaging session

Mice were placed head fixed in the functional imaging rig in front of a screen placed such as it covers the visual field of the right eye, contralateral to the craniotomy. Visual stimuli of the test block consisted of the presentation of one of two vertical sinewave gratings that drifted toward the right and were rotated clockwise by 45° and 135° (temporal frequency = 2 Hz, spatial frequency = 0.04 cycle per degree, contrast = 75%; duration: 3 s; intertrial interval: 3 s). Visual cues were presented in a pseudo-random order (30 to 60 presentations per block), such as the same stimulus could not be presented more than three times in a row. At the end of the imaging session, after a break of at least five minutes, we assessed the orientation tuning of the imaged neurons by presenting an orientation tuning block that consisted of the presentation of a series of drifting sinewave gratings (12 orientations evenly spaced by 30° and randomly permuted). The tuning block was placed after the test block not to interfere with the mouse performance. Indeed, trained mice tend to perform poorly after the presentation of hundred and twenty unrewarded trials. The spatiotemporal parameters of the orientation tuning stimuli were identical to those for the test block except for their duration (temporal frequency = 2 Hz, spatial frequency = 0.04 cycle per degree, contrast = 75%; duration: 1.5 s; intertrial interval: 3 s). All recording sessions contained an audiovisual block during which the two visual cues were paired with pure tones (45). Only the unimodal blocks were included in the analysis presented here (see Discussion). As scanning was not synced to the stimuli, a photodiode located at the stop left corner of the screen was used to detect the exact timing of the visual stimulus onset and offset. The photodiode signal was acquired along with the following signals: 1) a signal provided by the two-photon microscope, which indicated the onset of each frame, and 2) two analog signals encoding the orientation of the drifting grating. These signals were digitized (NiDAQ, National Instruments) and recorded with the software WinEDR (John Dempster, University of Strathclyde). Imaging sessions started by recording one thousand frames with the green and red channels. The red channel was used to exclude GABAergic neurons from the analysis.

### Behavioral training

After the naïve recording session, mice were water-deprived to 90% of their body weight and acclimated to head fixation on a spherical treadmill in custom-built, soundproof training rigs. Each rig was equipped with a monitor (Dell) and a water dispenser with a built-in lickometer (detecting licking using an infrared beam break). Data acquisition boards (National Instruments and Arduino) were used to actuate water delivery and vacuum after reward retrieval and record the animal licking. The monitor and data acquisition boards were connected to a computer that ran the custom-made training program scripted in MATLAB. Once animals reached the target weight and were acclimated to the training setup, they were trained to perform the orientation discrimination task. In this task, drifting sine-wave gratings oriented 45° below the vertical were paired with a water reward, and the animal was expected to lick (Go). Drifting gratings orthogonal to the Go signal (orientation 135°) signaled the absence of reward, and the animal was expected to withhold licking (No-Go) during those trials. When the stimulus instructed the animal to lick, the water delivery had to be triggered by the mouse licking during the third second of the stimulus presentation. No water was dispensed in the no-lick condition or if the mouse failed to trigger water delivery in the lick condition. If the animal responded correctly (Hit or Correct Rejection, CR), the intertrial interval was 3 s. If the animal responded incorrectly (Miss or False Alarm, FA), the intertrial interval was increased to 9.5 s as negative reinforcement. Performance was measured using the D’ statistic (D’=norminv(Hit rate) - norminv(False alarm rate), norminv = inverse of the normal cumulative distribution function). Animals are considered experts if their performance during training sessions is greater than 1.7 (probability of chance behavior < 0.1%, Monte Carlo simulation).

#### Trained and passive imaging sessions

Imaging sessions in trained mice (active or passive) were identical to the imaging sessions performed in naïve mice. However, in trained animals, imaging was performed while mice were performing the Go/No-Go discrimination task. In passive sessions, imaging was performed in trained mice at least 48 hours after water restriction was terminated. In this case, mice were not performing the task despite the water dispenser apparatus being present in front of the animal.

#### Timing of the response

The timing of the response (lick onset during Go trials and lick offset during NoGo trials) was quantified by averaging for each animal the lick traces recorded during the presentation of the Go and No-Go stimuli respectively. The mean trace across correct trials (hit and correct rejection respectively) was fitted by a sigmoid function. Lick onset (Go trials) and offset (NoGo trials) were defined as the inflection points of the rising (Go trials) or falling sigmoid fit (No-Go trials). The measure of lick cessation during NoGo trial was possible because mice presented a background licking activity that stopped shortly before the decision windows (3rd second of the stimulus presentation) during correct rejection trials.

### Data analysis

All the analyses detailed below were performed using custom MATLAB scripts.

#### Imaging data pre-processing

Calcium imaging frames were realigned offline to remove movement artifacts using the Scanbox algorithm (Neurolabware). A region of interest (ROI) was determined for each neuron using a semi-automatic segmentation routine. For every frame, the fluorescence level was averaged across the pixels of the ROI. Potential contamination of the soma fluorescence by the local neuropil was removed by subtracting the mean fluorescence of a 2-5 μm ring surrounding the neuron’s ROI, excluding the soma of neighboring neurons, and then adding the median value across time of the subtracted background. We then computed the fractional fluorescence from the background-subtracted fluorescence data. The fractional fluorescence (dF/F = (F – F0) / F0), was calculated with F0 defined as the median of the raw fluorescence measured during every inter-trial interval. The fractional fluorescence was then de-convolved into Action Potential related Events (APrE) using a benchmarked deconvolution algorithm (7, 18). Only neurons whose correlation between the inferred fractional fluorescence (convolved back from the inferred APrE) and the measured fractional fluorescence was greater than 0.8 were used for the analysis. This inclusion criterion removed neurons for which the calcium signal was too poor to allow the algorithm to accurately detect APrE. Trials were isolated and stored on a SQL database.

#### Tuning curves

The orientation tuning curve of each neuron was fitted to the APrE responses recorded during the tuning curve blocks using a Markov chain Mont Carlo sampling-based Bayesian method (14, 45) using a Poisson noise model. This algorithm fitted 4 different possible models to the observed responses (“circular Gaussian 360°” for orientation but not direction selective tuning curves, “circular Gaussian 180°” for strictly direction selective tuning curves, “direction selective circular Gaussian” for direction and orientation tuned neurons, and “constant” for untuned neurons). The fit that explained best the data was then selected. If the constant model was the best fit, the neuron was considered not tuned. The preferred orientation was defined as the peak of the fitted orientation tuning curve. When a neuron was not direction selective (i.e., responding equally to the same oriented stimulus moving in opposite directions), the preferred orientation was defined as the orientation included in the range [0 – 180]. The tuning width of the neurons (Fig. 5D,G) was computed from the fits of the tuning curves responses as the width of the fit, in degrees, at 50% of the peak value.

#### Orientation representation

To allow a full coverage of the orientation space, neurons were pooled across recording sessions based on the behavioral context of the imaging session (naïve, trained, passive). The preferred orientation of every neuron was assessed first (using the fits of the APrE responses recorded during the tuning block, see above). A relatively small contingent of neurons did not show significant orientation or direction tuning (naïve: 11.3%, n = 206/1830; trained: 16.7%, n = 357/2141). These neurons were not considered further for this type of analysis. For each stimulus of each group (e.g., 45° presented to trained mice), we generated an orientation space activity map of 1,000 resampled trials using the following procedure. For each resampled trial, 250 neurons (15% of the neurons of the database) were randomly selected, and for each neuron, one trial from the test block during which the stimulus of interest was presented (e.g., 45°) was selected at random. The activity of the neuron during the selected trial was included in an orientation*time matrix in which rows corresponded to the neurons’ preferred orientations (bin: 1°). The activity of neurons having the same preferred orientation was summed (instead of being averaged) to capture possible learning-induced changes in distribution of the preferred orientations between naïve and trained mice. Yet, because there is no change in the distribution of preferred orientation (see Results), the use of summation and average leads to identical results (not shown). The resulting trial matrix was baseline-subtracted and smoothed across the orientation space by a Gaussian weighted mean (std= 6°). The 1,000 resulting matrices were averaged to obtain the representation of the orientation of the stimulus by the V1 population. The flat preferred orientation procedure (used in Fig. 5B) was similar except that it included a rule forcing the selection of an equal number of neurons per orientation bin of 10°, flattening the distribution of the selected neurons’ preferred orientation. To determine the distribution across trials of the peak of activity, the orientation space activity map for each resampled trial was averaged across the first second of the stimulus presentation. The peak of activity in the orientation space was then determined for each resampled trial and the distribution of the 1,000 resulting orientations was fitted by a mixture of von Mises distributions (two to four mixture components) using mvmdist (https://github.com/chrschy/mvmdist), a Matlab package for probabilistic modeling of circular data. We made sure that our findings did not depend on the number of neurons used in the resampling procedure (250) by reproducing the results presented here using 1,000 samples of 500 and 1,000 neurons randomly selected in the databases (not shown).

#### Locomotion and pupil size

Locomotion (ball motion) and pupil size traces were lowpass filtered (0.25Hz) and then z-scored for each session. For pupil size, a small number of trials were discarded because either the mouse was blinking or the position of the eye was not allowing us to reliably assess the size of the pupil. To reflect the resampling procedure used to determine the representation of the orientation of the visual stimuli by the V1 neuronal population (see above), we performed the same resampling procedure, replacing the activity of the neurons by the amount of locomotion or pupil size data. To evaluate the impact of the differences of locomotion and pupil size between groups, we sorted all trials in 4 categories according to the amount of locomotion/pupil size. The cut-off values were based on the quartile values of the trained group, as it was the group displaying the largest amplitude of locomotion and greater pupil size values compared to the other groups.

#### Mean tuning curves comparison

To compare the tuning curves of neurons in specific bins of orientation preference (Fig. 7I,J), the tuning curve fit of all considered neurons was circularly shifted so that the peak would be centered at 0°. To visualize the data underlying the averaged fits, the APrE responses were shifted by the same number of degrees. For example, the data from a neuron with a preferred orientation of 45° would be shifted such that its response to 0° would show at −45° on the plot, its response to 30° would show at −15°, etc. The median of each individual neuron’s responses was then computed, and for each degree of the orientation space the medians from all neurons falling in that degree (that were equally shifted) were averaged.

#### Orientation Selectivity Index (OSI)

Individual OSI were calculated from the fitted tuning curve, following the formula OSI =(R_pref_ + R_opp_ - (R_orth+_ + R_orth-_))/(R_pref_ + R_opp_) (44), where R_pref_ stands for the tuning curve amplitude in APrE/trial at the preferred direction, R_opp_ that of the opposite direction, and R_orth +/−_ stands for the tuning curve amplitude at the orthogonal orientation. The probability distributions of OSI were binned with a width of 0.01 and smoothed with a moving average of 10 points. The confidence intervals of the distributions were obtained by bootstrapping the OSI values 10000 times and taking the .025 and .975 percentiles of each bin.

#### Responsive neurons

Responsive neurons were identified using a binomial test that compared the probability of having at least one APrE during the stimulus presentation, to the probability of having at least one APrE during the pre-stimulus time. We first estimated the probability of each neuron to fire APrE during the last second of the intertrial interval (binofit function (MATLAB, MathWorks); Clopper–Pearson method). Then, we compared the probability of observing activity during a one second window sliding across the presentation of the stimulus to the upper bound of the 99% confidence interval of the probability of activity calculated for the intertrial interval. A neuron was considered responsive if the probability of APrE in at least one of the sliding windows was greater than the confidence interval intertrial activity probability. The use of a binomial model to detect neural responses was motivated by the sparseness of the observed activity. Indeed, 84% of the intertrial interval windows did not contain any APrE; only 8% contained a single APrE and another 8% two or more. Hence, assessing the presence of any amount of APrE captured most of the variability of the neuronal activity. The use of a Wilcoxon test approach to compare either the responsiveness (1s pre-stimulus vs post-stimulus) or the selectivity (1s post-stimulus 45° vs 135°; see (53)) led to similar results and conclusions (not shown).

### Statistics

#### Permutation tests

To determine if the means across trials of two pools were significantly different, we compared the value obtained to the distribution of 1,000 or 10,000 differences obtained when the pool labels were shuffled. The two-tailed confidence interval of the null hypothesis at the alpha level 0.05 was defined as the 2.5 and 97.5 percentile of the distribution obtain from the permutations. The difference between the observed means was considered significant if located outside the confidence interval of the null distribution.

Comparison of preferred orientation distributions. Comparison of the preferred orientation distribution was achieved with an Anderson-Darling nonparametric test, adjusted for ties (36). Circular statistics. Circular statistics were computed with the Circular Statistics Toolbox for MATLAB (5). The difference between groups was considered significant if the values of the confidence interval boundaries did not include zero.

#### Neuronal activity

The significance of the difference in activity between naïve and trained mice, for neurons belonging to a specific neuronal group defined by their orientation preference, was assessed by a two-sample Wilcoxon rank sum test with a significance threshold of p < 0.05. Effect size.

Because large number of observations can lead to conclude on significant differences of even irrelevantly small magnitude, we complimented the tests with an effect size measure, Hedges’g, using a MATLAB toolbox (31). Traditionally, values under 0.2 correspond to the absence of effect, values between 0.2 and 0.4 are considered to reflect a small effect, values between 0.4 and 0.6 correspond to a medium effect, and values larger than 0.6 correspond to a large effect size.

## Results

To evaluate the changes induced by learning on the representation of oriented stimuli in V1, we presented the same sets of drifting gratings to naïve mice (14 mice and 21 sessions) and to mice trained to use those cues to perform an orientation discrimination task (trained mice, 10 mice and 16 sessions). V1 neuronal activity was recorded by calcium imaging in mice head-fixed on a spherical treadmill (45) (**Fig. 1A**). Recording sessions consisted first of a test block during which two orthogonal drifting gratings were presented (45° and 135°; **Fig. 1B**). Trained mice were taught to discriminate between the Go cue (45° drifting grating) and the No-Go cue (135° drifting grating; **Fig. 1C**). A task trial consisted in the presentation of one of the two test stimuli for 3 seconds. Trained mice had to respond to the test cues during the third second of the trial (**Fig. 1D**). As locomotor activity and arousal modulate the neuronal activity in V1 (52, 65), treadmill movement and pupil size were recorded simultaneously with the calcium signal and the licking activity (**Fig. 1E,F**). The calcium signal was deconvolved into Action Potential related Events (APrE) using a benchmarked algorithm (18) (**Fig. 1F**). At the end of the recording session, we presented a tuning block consisting of gratings of six different orientations (distinct from the two orientations of the test block) drifting in the two opposite directions (**Fig.1G**). Note that for clarity, we will use the term “orientation” to describe both the orientation and the direction of the drifting gratings, therefore orientations will be ranging from 0 to 360°. The neuronal responses to the tuning block were fitted using a resampling-based Bayesian method to determine the tuning curve and preferred orientation of each recorded neuron (14, 45) (**Fig. 1H,I**). When neurons were orientation selective but not direction selective (see example in **Fig. 1H**), the preferred orientation was assigned to the sector 0° - 179°. As a result, the distribution of preferred orientations across the population was inflated in that half of the possible orientation values (**Fig. 1J**). During the behavioral task, trained mice successfully licked during the reward period for Go trials and refrained from licking during No-Go trials (**Fig. 2A**). They performed the task with a hit rate of 94 ± 8% and a correct rejection rate of 67 ± 13% (median ± median absolute deviation (m.a.d.); p=9.8 x 10-5; Kruskal-Wallis Test: χ^2^ = 15.2; **Fig. 2B**). Altogether, the mean sensitivity index (D’) across all the sessions was 2.1 ± 0.2 (16 sessions in 10 mice, **Fig. 2C**).

**Fig. 1.**
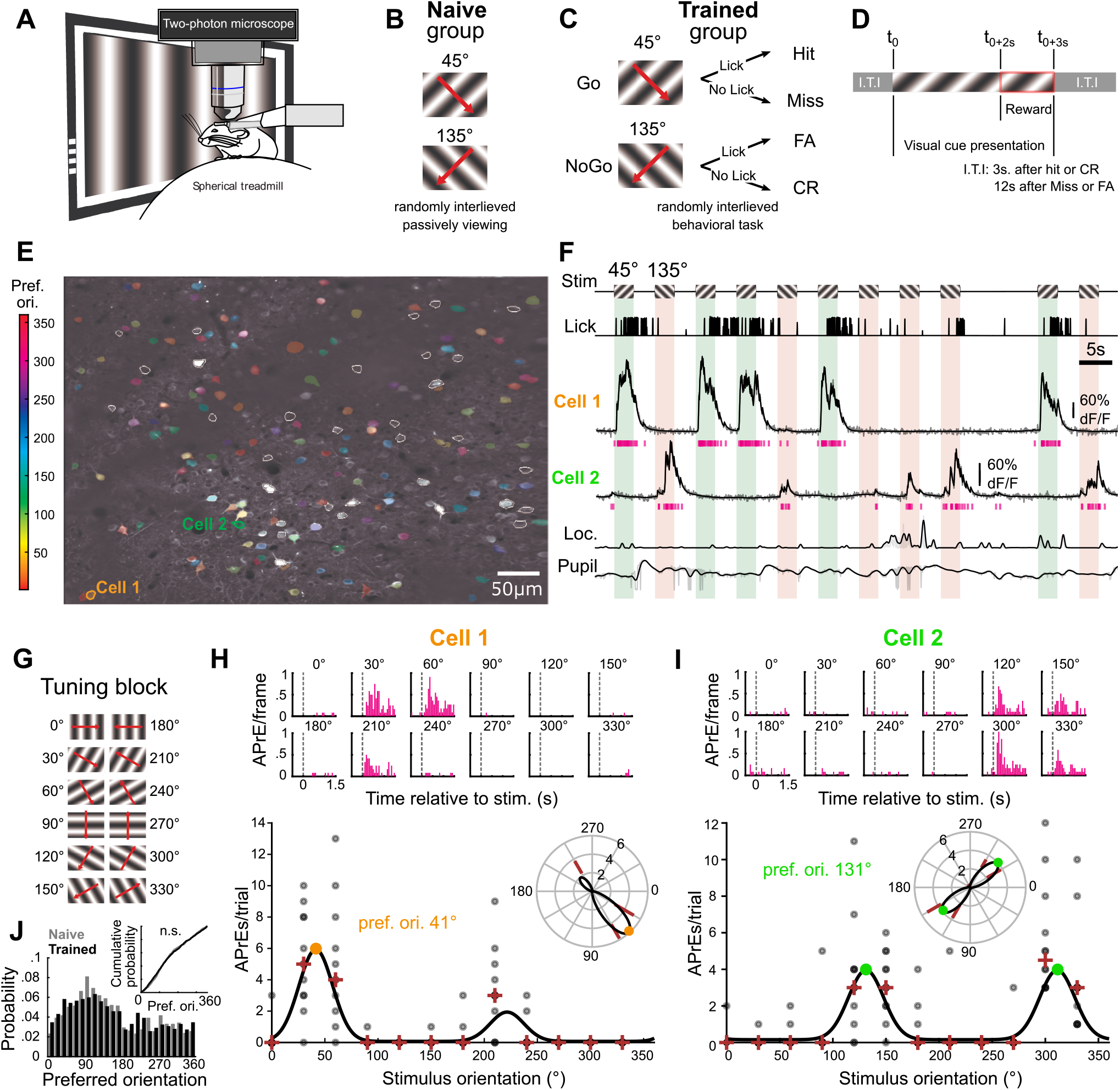
Experimental design. **A**. Imaging setup. **B**. Test block for the Naïve group. **C**. Schematic description of the Go/No-Go task for the Trained group. FA = False Alarm; CR = Correct Rejection. **D**. Temporal structure of a task trial. I.T.I.: intertrial interval. **E**. Representative two-photon image of V1 layer 2/3. Contours indicate segmented neurons. Colors refers to the neurons’ preferred orientation. White contours indicate untuned neurons. **F**. Example segment of recording during the task. From top to bottom: visual stimulus, mouse licking activity, fractional fluorescence (dF/F; grey traces) of the two cells shown in (E) with the deconvolved Action Potential related Events (APrE, pink) and the reconvolved signal (black), mouse locomotion, and mouse pupil size. **G**. 12 orientations presented during the tuning block. **H-I**. Tuning curves of the neuron presented in E-F. Top: Mean APrE response to the 12 orientations. Bottom: Fitted tuning curve (black), APrE count per trial (gray circles), median count across trial (red cross). Inset: same data wrapped around polar coordinates. Colored dots indicate the preferred orientation. **J**. Preferred orientation distribution of neurons recorded in naïve and trained mice. Inset: Cumulative probability of the two distributions. n.s.: non-significant, Anderson-darling test, p=0.70.

**Fig. 2.**
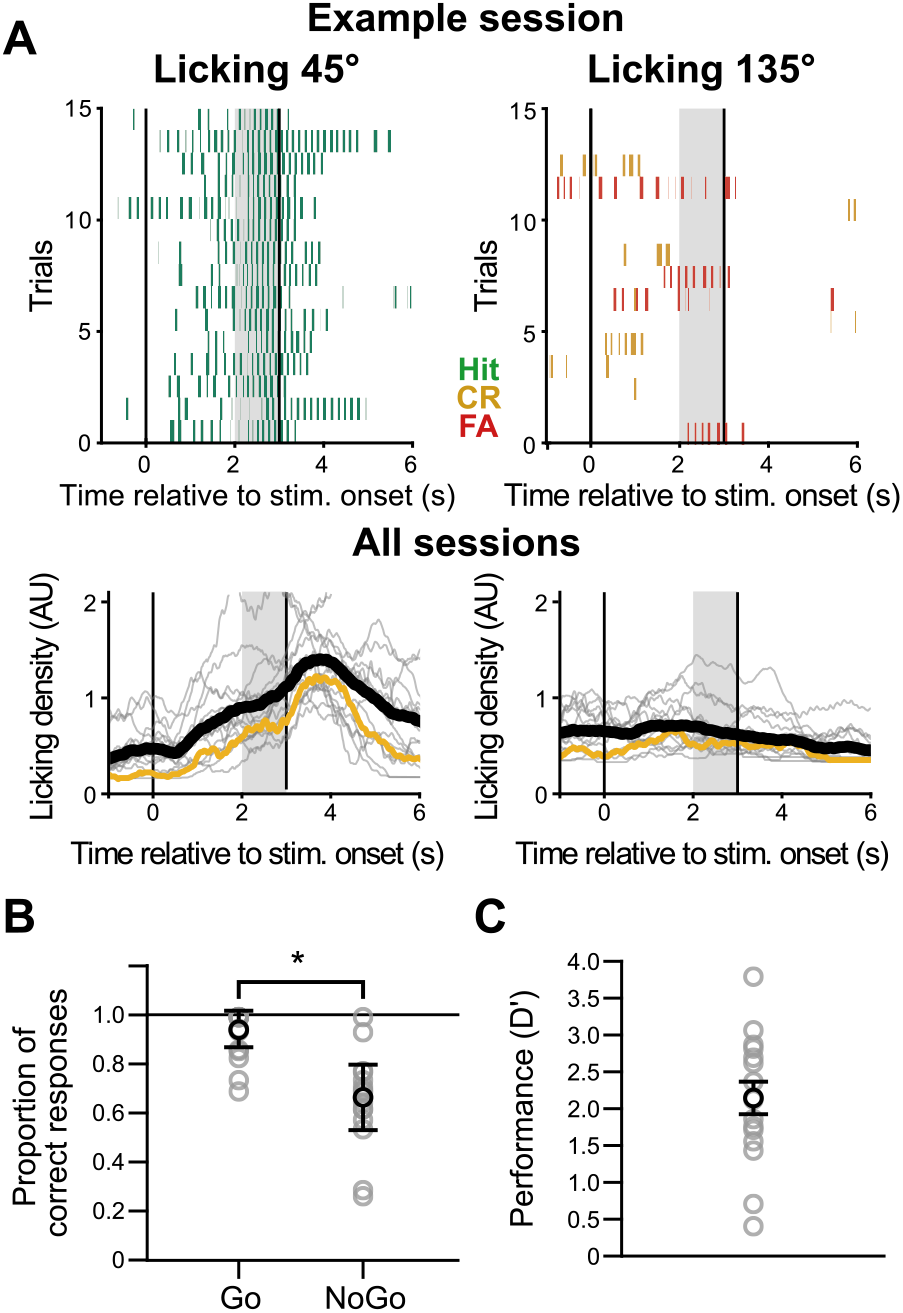
Behavioral performance of the trained mice. **A**. Top: example of licking responses during the presentation of a 45°oriented grating (left) and 135°oriented grating (right). From the same session as depicted in Fig. 1F. Bottom: Mean licking density within sessions (grey line) and across sessions (black line). The session used as an example above is highlighted in yellow. Grey shaded areas indicate the behavioral response window. **B**. Proportion of correct responses to the presentation of the Go and No-Go cues for the ‘trained’ group. * p = 0.0001; χ^2^ = 15.2; n = 16 sessions in 10 mice. **C**. Performance (D’) of the trained group for each recording session.

### Sharpened representations in the orientation space after training

To determine how the visual stimuli were represented in the orientation space of V1, we generated an orientation space activity map for each test stimulus using a re-sampling approach (see Material and Methods). This map captured the density of APrE evoked during the stimulus presentation as a function of the preferred orientations of the recorded neurons (**Fig. 3A**). We restricted our analysis in time by averaging the orientation space activity map across the first second of the visual stimulus presentation, i.e., before the licking response of trained mice (Go response time, mean ± s.e.m.: 1.8 ± 0.1 s, No-Go response time 1.9 ± 0.1 s, 16 sessions in 10 mice). The resulting profile of activation was broader in naïve (**Fig. 3B**) than in trained mice (**Fig. 3C**), which meant that the test stimuli in naïve mice were activating neurons across a larger domain of the preferred orientation space. The sharpening found in trained mice was due to the suppression of the activity of neurons tuned for orientations flanking the orientation of the test stimulus (**Fig. 3D,E**). Indeed, the responses of neurons whose preferred orientation flanked the orientation of the test block cue (15° wide bins centered 15° away from the test stimulus orientations) were significantly smaller in trained mice than in naïve mice (**Fig. 3F**, “flank” group, For 45°: n_naive_ = 150 neurons; n_trained_ = 196; Wilcoxon rank sum test: p = 0.002, Hedges’ g for effect size: g = 0.55; For 135°: n_naive_ = 207 neurons; n_trained_ = 183; p = 0.03, g = 0.33). Meanwhile, no significant difference was found between the activity of neurons of naïve and trained mice tuned to the orientation of the test stimuli (“tuned” group, 15° bins centered on 45° and 135°, **Fig. 3F**,

**Fig. 3.**
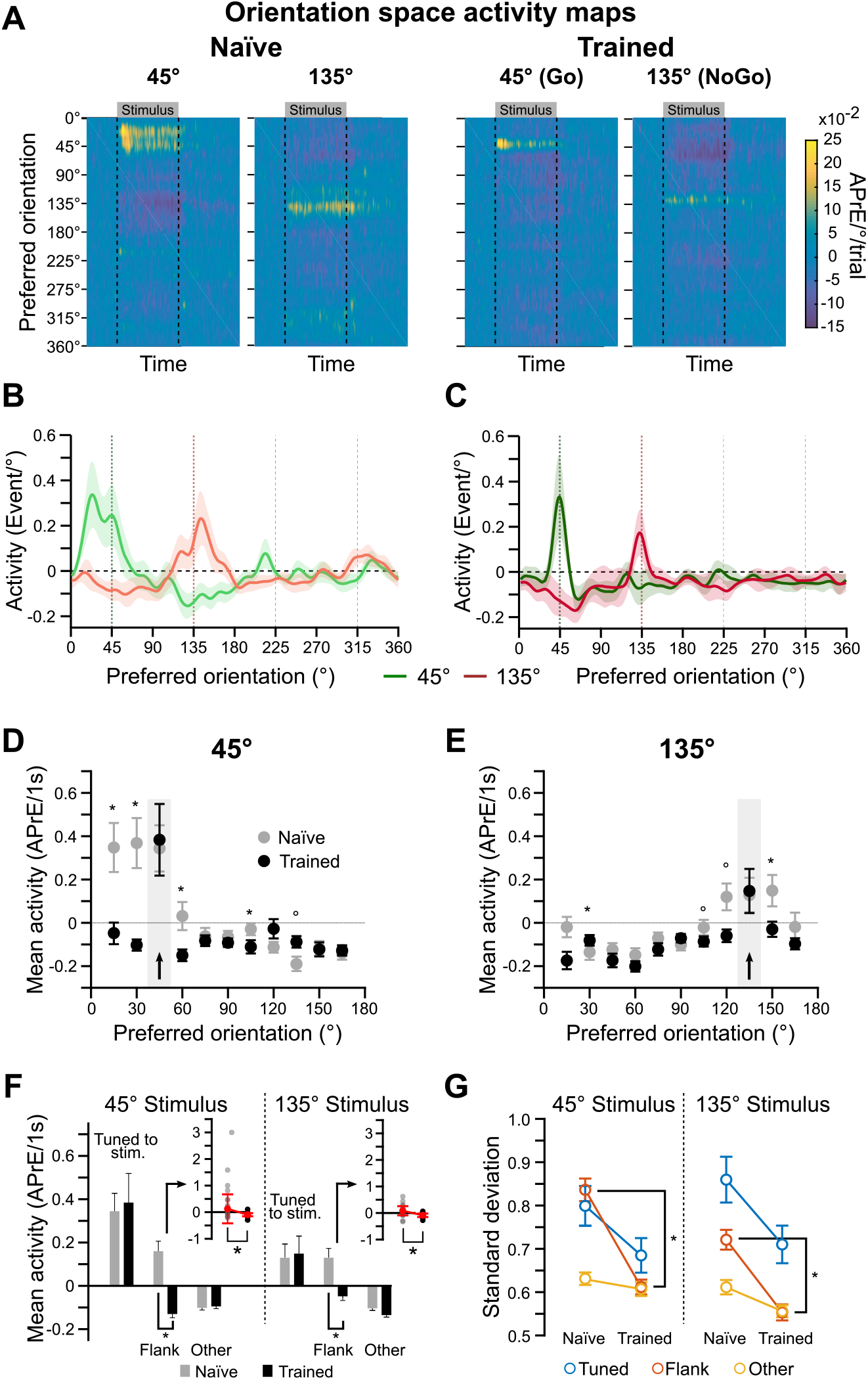
Representations of the test stimuli in the V1 of naïve and trained mice. **A**. Map of the V1 activity evoked in the orientation space by the presentation of the test stimuli (45°and 135°) in naïve (left panels) and trained mice (right panels). **B**. Average of the orientation space activity map shown in (A) over the first second of the presentation of the 45°(green trace) and 135° (red) test stimuli in naïve mice. Shaded areas indicate the standard deviation across time. **C**. Same representation as in (B) in trained mice. **D**. Average across V1 neurons (mean ± s.e.m.) of the mean response relative to baseline (in APrE.s-1) in response to the presentation of the 45° test stimulus in naïve (grey) and trained (black) mice. Neurons are grouped by preferred orientation (bin: x-axis value ± 7.5°). * p < 0.05, o p<0.01. Two-sample Kolmogorov-Smirnov test. **E**. Same representation as in (A) for the neuronal responses to the presentation of the 135°test stimulus. **F**. Responses to the presentation of the 45°and 135°stimuli of the neurons tuned for the test stimuli (Tuned: 45°or 135°±7.5°), neurons tuned for orientations flanking the test stimulus orientation (Flank for 45°: 30°±7.5°and 60°±7.5°; Flank for 135°: 120°±7.5° and 150°±7.5°), and the other neurons between 0° and 180°. *: p < 0.05, Wilcoxon rank sum test. Inset: same analysis performed for each session (grey dots) and across session average (in red, median ± m.a.d.). **G**. Standard deviation of the neuronal response across trials for the same groups. *: p < 0.05, Wilcoxon rank sum test.

For 45°: n_naive_ = 73 neurons; n_trained_ = 101; Wilcoxon rank sum test: p = 0.55; For 135°: n_naive_ = 89 neurons; n_trained_ = 95; Wilcoxon rank sum test: p = 0.49). Neurons tuned to other orientations (i.e., not belonging to the tuned or flank group, and belonging to the 1-180° sector) did not show a significant modulation between naïve and trained animals (Fig. 3F, “other” group, For 45°: n_naive_ = 766 neurons; n_trained_ = 739; Wilcoxon rank sum test: p = 0. 87, Hedges’ g for effect size = 0.06; For 135°: n_naive_ = 693 neurons; n_trained_ = 758; p = 0.01, g = 0.14 i.e.., no effect, see Methods). This suppression of flanking neurons in trained mice was consistent across recording sessions (**Fig. 3F** insets) and associated with a significant decrease of the response variance across trials (**Fig. 3G**, For 45°: Wilcoxon rank sum test: p = 4×10-5; For 135°: Wilcoxon rank sum test: p = 4×10-4). We therefore tested the hypothesis that the sharper representation found in trained mice resulted from trial-to-trial responses more stable in the orientation space than in naïve mice. To do so, we determined where the evoked activity profiles peaked in the orientation space for each resampled trial (**Fig. 4A-D**). The resultant distributions were fitted with von Mises distributions. The widths of those fits (σ) were used as indicators of representation stability across trials, as more stable representations in the orientation space would be characterized by narrower distributions. We found the fit width to be significantly broader in naïve mice (σ45° = 13.2°; C.I._95%_: [12.5 – 13.8]; σ135° = 19.2°, C.I._95%_: [18.2 – 20.1]; **Fig. 4E**) than in trained mice (σ45° = 2.5°, C.I._95%_: [2.4 – 2.7]; σ135° = 2.5°, C.I._95%_: [2.4 – 2.7]; Fig. 4F). We ruled out the hypothesis that this result could be caused by a single outlier session using a jackknife analysis, repeating the measure of the fits’ width but leaving one session out (**Fig. 4G**). Thus, the representations of task-relevant cues in the orientation space were more stable and accurate in trained animals, with neurons preferring the orientation of the test cues reliably being the most active across trials.

**Fig. 4.**
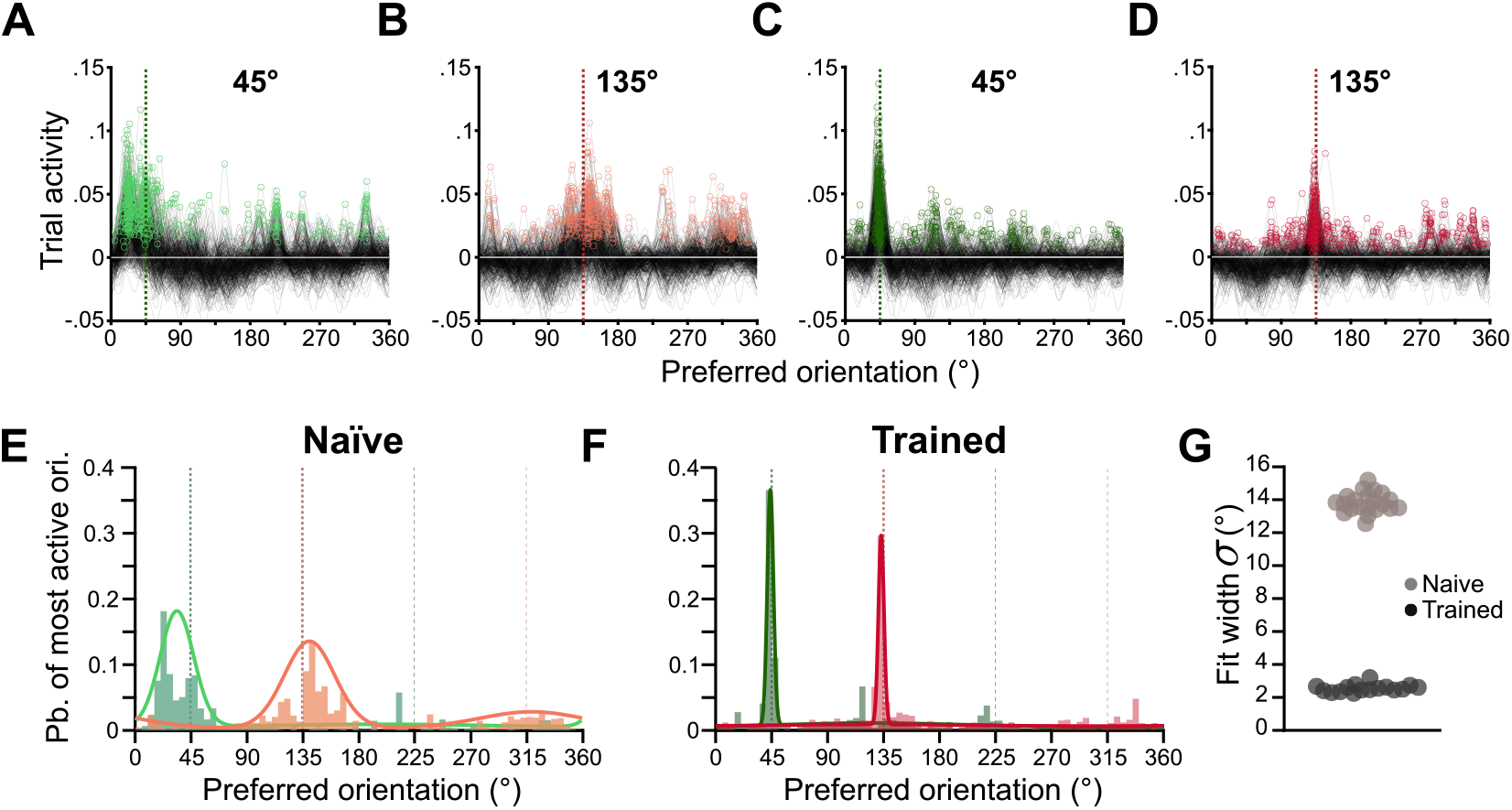
Stability of the test stimulus representations in naïve and trained mice. **A**. Superimposition of 500 response profiles (average of the orientation space activity map over the first second of the stimulus presentation as in Fig. 3B,C but for individual resampled trials) in response to the presentation of the 45°test stimulus in naïve mice. Colored dots indicate the peaks of activity for the individual response profiles. **B**. Same as in (A), for the 135°test stimulus in naive mice. **C-D**. Same as A-B for trained mice. **E**. Probability distribution in the orientation space of the peak of activity of the resampled trials for the presentation of the 45°(green) and 135°(red) test stimuli in naïve mice. The peak distributions were fitted with a two component von Mises mixture model. **F**. Same as in (E). for trained mice. **G**. Width of the von Mises distribution fitted on the data obtained through a leave-one-out jackknife procedure.

The sharpening of the representation of the test stimuli in trained mice could not be explained by a difference in the distribution of preferred orientations (those distributions were not different, Anderson-Darling non-parametric test: p=0.70; **Fig. 1J**). Moreover, changing the neuron selection procedure (such as the randomly selected neurons formed a flat distribution of preferred orientations) did not change the general profiles of the evoked activity in the orientation space (**Fig. 5A**). Because the behavioral state and movements of the animals can modulate the activity of V1 neurons (22, 50, 52, 63, 65), we analyzed the representations as a function of the amount of licking, locomotion and pupil size.To assess whether licking in the first second could contribute to the cue representation sharpening in trained animals, we sorted trials according to the presence or absence of licking activity (**Fig. 5B**). Using trials with and without early licking lead to similar activity profiles in the orientation space (**Fig. 5C**), as shown by the similar von Mises fits of the peak distributions that were comparable for the 45° stimulus (σ45°_NoLick_ = 2.75°; C.I._95%_: [2.63 – 2.9]; σ45°_Lick_ = 2.73°; C.I._95%_: [2.6 – 2.87]], **Fig. 5D**) and slightly wider for the 135° stimulus in the absence of licking (σ135°_NoLick_ = 3.91°; C.I._95%_: [3.72 – 4.12]; σ135°_Lick_ = 3.39°; C.I._95%_: [3.23 – 3.57], **Fig. 5D**). Most importantly, with or without licking, the width of the fits stayed largely different from that observed for the naïve group (**Fig. 5E**). Trained animals displayed a greater amount of locomotion during baseline and increased their locomotor activity during the presentation of the test stimuli. The locomotor activity also was the largest during rewarded trials (**Fig. 5F**). Pupil size showed a similar pattern, with a greater pupil dilatation overall in trained mice (**Fig. 5G**). To determine whether differences in arousal and locomotion could explain the difference in orientation representations observed between naïve and trained animals, we measured the width of the peak distributions after sorting the trials according to the amount of locomotion or pupil dilatation measured during the first second post stimulus onset (**Fig. 5F,G**). The distribution widths remained well separated between naïve and trained mice regardless of the amount of locomotion or pupil dilatation. Even when there was no locomotion (Quartile 1, mean locomotion value = 0), the stability in the orientation space of the response in the trained group was greater than in the naïve group, excluding a role of locomotion and arousal in our results. Finally, we compared the quality of the tuning curve fits of naïve and trained mice. Indeed, noisier responses during the tuning curve block would decrease the accuracy of the neurons’ preferred orientation estimation, and more loosely categorized neurons could in turn create a broader activity profile in the orientation space. We therefore computed the standard deviation of the responses to the presentation of orientations eliciting the most activity for each neuron and did not find a significant difference between the naïve and trained groups (naïve: median = 0.90, 1st and 3rd quartiles = [0.30, 1.50], trained: 0.86, [0.39, 1.33], permutation test, p=0.03, effect size Hedges’ g=−8.1×10-2 **Fig. 5H,I**). We also compared the mean squared error (MSE) between neuronal responses and the fitted tuning curves in naïve and active animals (**Fig. 5H,J**). No significant difference was found for the entire recorded populations (naïve: 0.69, [0.36, 1.23]; trained: 0.67, [0.33, 1.21], p=0.21, g=4.4×10-2), nor for the neurons tuned to the test orientations (naïve: 0.71, [0.32, 1.21]; trained: 0.60, [0.29, 1.07], p=0.14, g=−2.5×10-2) and flanking orientations (naïve: 0.63, [0.33, 1.23]; trained: 0.62, [0.31, 1.09], p=0.14, g=−2.5×10-2). Finally, we compared the width of the tuning curve in trained and naïve mice as flatter tuning curves could potentially lead to a noisier estimate of the preferred orientation of the neurons. The tuning width was similar for neurons tuned to the test orientations (naïve: 35°, [22°, 42°]; trained: 35.5°, [26°, 43°], p=0.58, g=0.16), for neurons tuned to the flanking orientations (naïve: 32°, [22°, 43°]; trained: 33°, [23°, 42°], p=0.89, g=2.0×10-2), and for neurons tuned for all other orientations (naïve: 31°, [22°, 42°]; trained: 32°, [22°, 44°], p=0.96, g=0.04, **Fig. 5H,K**).

**Fig. 5.**
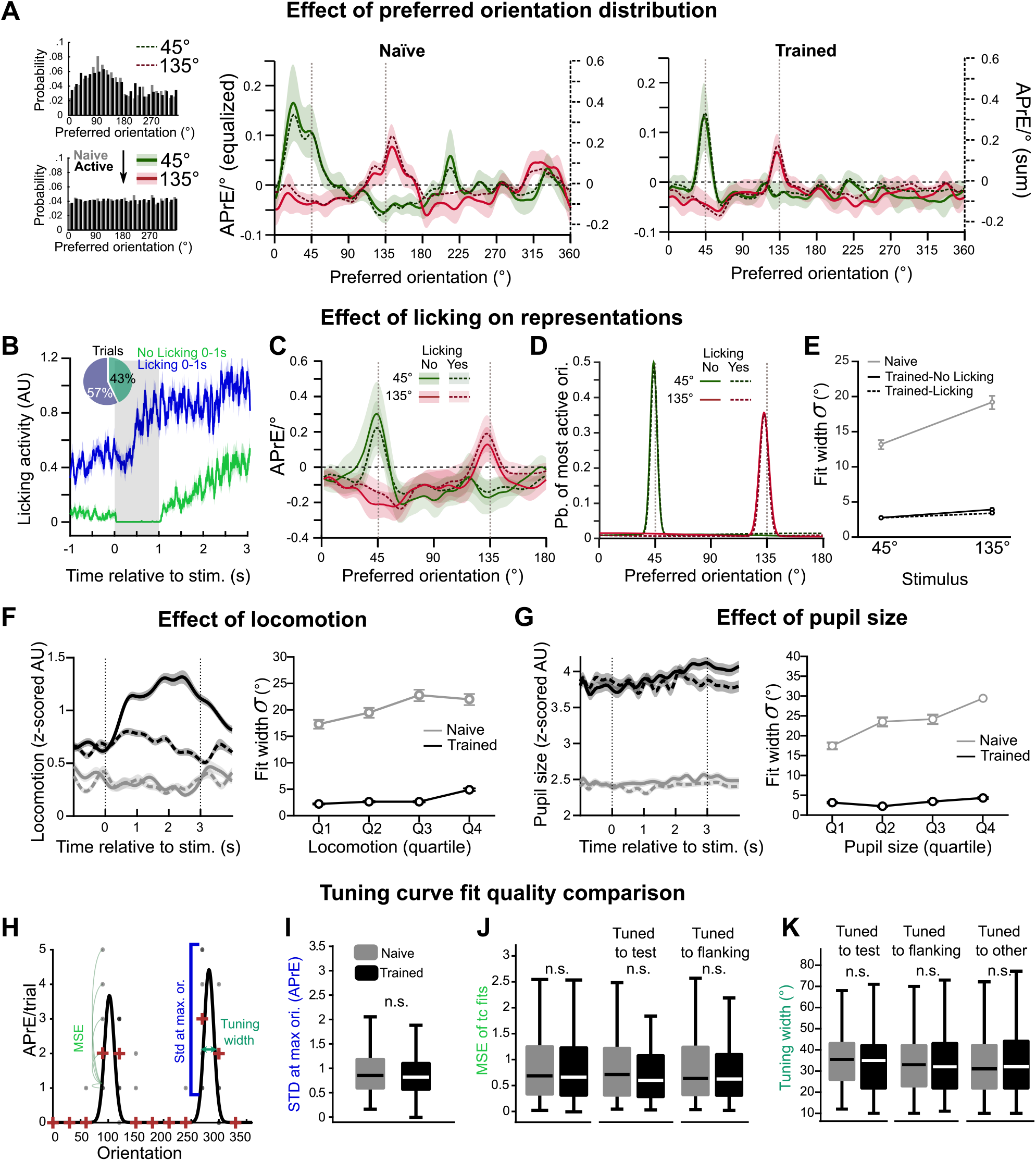
Impact of preferred orientation distribution, locomotion, arousal, and tuning curve fits on the representation of the stimulus orientation in V1. **A**. Top left panel: observed preferred orientation distribution. Bottom left panel: distribution obtained after modifying the resampling approach to use the same number of neurons across the orientation space. Center and right panels: activation profile over the first second of the presentation of the 45°(green) and 135°(red) test stimuli in naïve (center panel) and trained (right panel) mice when the sorting of neurons is adjusted for preferred orientation. Dotted lines (and right y-axis): profiles (same as in **Fig. 3B**) preserving the observed distribution of preferred orientations. **B**. Mean licking activity in arbitrary units (A.U.) for all the test trials (Go and NoGo) with (blue) or without (green) licking activity in the first second of stimulus presentation. Shaded areas indicate the standard deviation. Inset: proportion of trials with and without early licking. **C**. Activation profiles in the orientation space over the first second of the presentation of the 45°(green) and 135°(red) test stimuli for trials with (dashed lines) and without (plain lines) licking, for the trained animals. Shaded areas indicate the standard deviation. **D**. Two component von Mises mixture model fits of the resampled trials peak distribution, with (dashed lines) and without (plain lines) licking. **E**. Fit width σ ± C.I.95% for 45°and 135°stimuli responses of trained animals (black) for trials with (dashed line) and without licking (plain line), and of naïve animals (grey). **F**. Left panel. Mean locomotion time course (in z-scores) during the presentation of 45°(continuous line) and 135°(dotted lines) cues in naïve and trained animals. The traces were obtained using the same resampling method as for the neuronal representations. Mean ± standard deviation. Right panel. Width of the peak distribution across trials as a function of the quantity of locomotion occurring during the first second after stimulus onset. All trials were sorted according to the amount of locomotion observed during the first second of the trial, with quartiles cutoff values computed from the trials in the trained group. σ ± C.I.95%. **G**. Same as in (F) but for pupil size. **H**. Tuning curve (see Fig.1H) of a neuron of the dataset with a schematic representation of the MSE (mean squared error), the standard deviation at max. orientation and tuning width. **I**. Standard deviation at max. orientation in naïve and trained mice. **J**. MSE of the tuning curve fits in naïve and trained mice for all the neurons (left), neurons tuned to the test stimulus orientation (45 and 135 ± 7.5°; center) and neurons tuned to orientations flanking the orientation of the test stimulus in naïve and trained mice (30, 60, 120 & 150 ± 7.5°). **K**. Width of the tuning curves of neurons tuned to the test stimulus orientation, neurons tuned to orientations flanking the orientation of the test stimulus, and neurons tuned to all other orientations, in naïve and trained mice. n.s. indicates a non-significant permutation test p-value and Hedges’ g effect size.

### Distortion of the orientation space in trained mice

We wondered if the modulation of the representation of the test stimuli in the orientation space was specific of the task orientations (as it would be expected if task training was inducing local neuronal plasticity), or if similar changes would be also present for orientations other than those of the test block (as it would be expected if they were the results of a non-specific neuronal modulation associated to the behavioral context; (22, 52, 65)). We therefore generated orientation space activity maps for the twelve stimuli of the tuning block in naïve and trained mice. In trained mice, these maps presented unexpected distortions when orientations flanking the Go and NoGo stimuli were presented (30° and 60°, 120° and 150°, see the 60° representation in **Fig. 6A,B**, and all the orientations in **Fig. 6C**). Those distortions were characterized by a peak of activity of the resampled trials consistently located at the orientations of the test stimuli instead of being found at the orientation of the presented stimulus. Stimuli opposite to those flanking orientations (210°, 240°, 300° and 330°) also evoked misplaced representations on the 1-180° sector (**Fig. 6C**). To characterize this distortion of stimulus representation in the orientation space, we compiled the distributions of the resampled trial activity peaks for all the flanking orientations (30° and 60°, 120° and 150°, **Fig. 7A**). Note that the distributions of the 60° and 150° stimuli were mirrored to allow for the superimposition). The resulting distribution was fitted by a mixed model of von Mises distributions whose best component (most variance explained) was tightly centered at the orientation of the test stimulus (peak location relative to the presented orientation ± CI95%,: +15.5 ± 0.7°; σ = 3.9°, σ C.I._95%_ : [3.7 – 4.1]) while the second largest proportion was broader (σ = 10.9°, σ C.I._95%_ : [10.3 – 11.4]) and away from the orientation of the tuning block stimulus (peak location relative to the presented orientation: −9.4 ± 2.0°; **Fig. 7A**). This deformation was not found with stimuli distant to the test stimuli (0°, 90° 180°, and 270°; peak location relative to the presented orientation: + 2.6 ± 1.7°; **Fig. 7B**) even though the width of the fit was significantly smaller (σtrained = 9.5°, σ C.I._95%_ : [9.0 – 10.0] vs. σnaive = 13.8°, σ C.I.5%: [13.1 – 14.5]). Finally, the peak of activity in the orientation space for tuning block stimuli opposite to test stimuli was centered (210°, 240° 300°, and 330°; peak location relative to the presented orientation: −2.3 ± 3.3°) and slightly broader after training (σtrained = 17.4°, σ C.I._95%_: [16.6 – 18.3] vs. σnaive = 15.6°, σ C.I._95%_: [14.9 – 16.4]; **Fig. 7C**). In naive mice, the peak of activity of the resampled trials was consistently located at the orientations of the presented stimulus for flanking (**Fig. 7D**), distant (**Fig. 7E**) and opposite orientations (**Fig. 7F**). We did not find significant differences between the width of the peak distribution for the flanking (σ = 14.6°, C.I._95%_: [13.8 – 15.3]), distant (σ = 13.8°, C.I._95%_: [13.1 – 14.5]), or opposite stimuli (σ = 15.6°, C.I._95%_: [14.9 – 16.4]; **Fig. 7D-F**). These results suggest that the repeated presentation of the 45° and 135° stimuli during a session does not impact the representation of flanking orientations unless the stimuli acquire behavioral significance through learning. As a result of this distortion, orientations close to the task-relevant cues seem to be represented in trained animals as the task-relevant cues themselves. These results, coupled with the sharpening observed in the representation of the test stimuli, suggested the presence of a filter applied to the orientation space through the specific suppression of the neurons preferring the orientations flanking the orientation of the learned cues. To visualize this filter, we sorted the neurons by preferred orientation (bins of 7.5°) and compared their mean response to the presentation of the tuning block stimuli flanking the test stimuli on the left (i.e. 30° and 120°, **Fig. 7G** left panels). We then subtracted those two curves to determine how each group of neurons was modulated by the training (**Fig. 7G**, right panel). The responses of the neurons tuned for the orientation of the presented stimulus (30° and 120°) were significantly suppressed in trained mice compared to naïve mice (**Fig. 7G**) while the responses of the neurons tuned for the closest test stimulus (45° or 135°) were similar (**Fig. 7G**). The same result was obtained when performing the same analysis for the presentation of the tuning block stimuli flanking the test stimuli on the right (i.e. 60° and 150°; **Fig. 7H**). The fact that flank neurons were suppressed during the tuning block such that they were not the most activated neuronal population during the presentation of their own preferred stimulus could question their classification as neurons preferring these orientations. We therefore centered and then averaged the tuning curves of neurons tuned for the test stimuli (45° or 135° ± 7.5°; **Fig. 7I** top left inset) and the flanking stimuli (30°, 60°, 120° and 150° ± 7.5°, Fig. **7J** **top left inset**) in naïve and trained mice. While the tuning curves of test neurons were similar in naïve and trained mice (mean peak amplitude in APrE/trial ± CI95%, naïve: 0.80 ± 0.14, trained: 0.81 ± 0.21, permutation test p = 0.56, Hedges’ g = −0.01, **Fig. 7I**), the amplitude of the tuning curves of flank neurons was smaller in trained than in naïve mice (mean peak amplitude in APrE/trial ± CI95%, naïve: 0.83 ± 0.11, trained: 0.50 ± 0.051, p = 0, g = 0.39, **Fig. 7J**). This change in mean tuning curve amplitude was reflected in the Orientation Selectivity Indexes (OSI) distributions. Indeed, the distributions were comparable for neurons tuned to the test stimuli between naïve and trained animals (**Fig. 7I** top right inset) but for flanking neurons we observed a decrease in the proportion of highly selective neurons in trained animals (**Fig. 7J** top right inset, OSI>0.9) coupled with an increase in the proportion of medium OSI values (0.5 to 0.7).

**Fig. 6.**
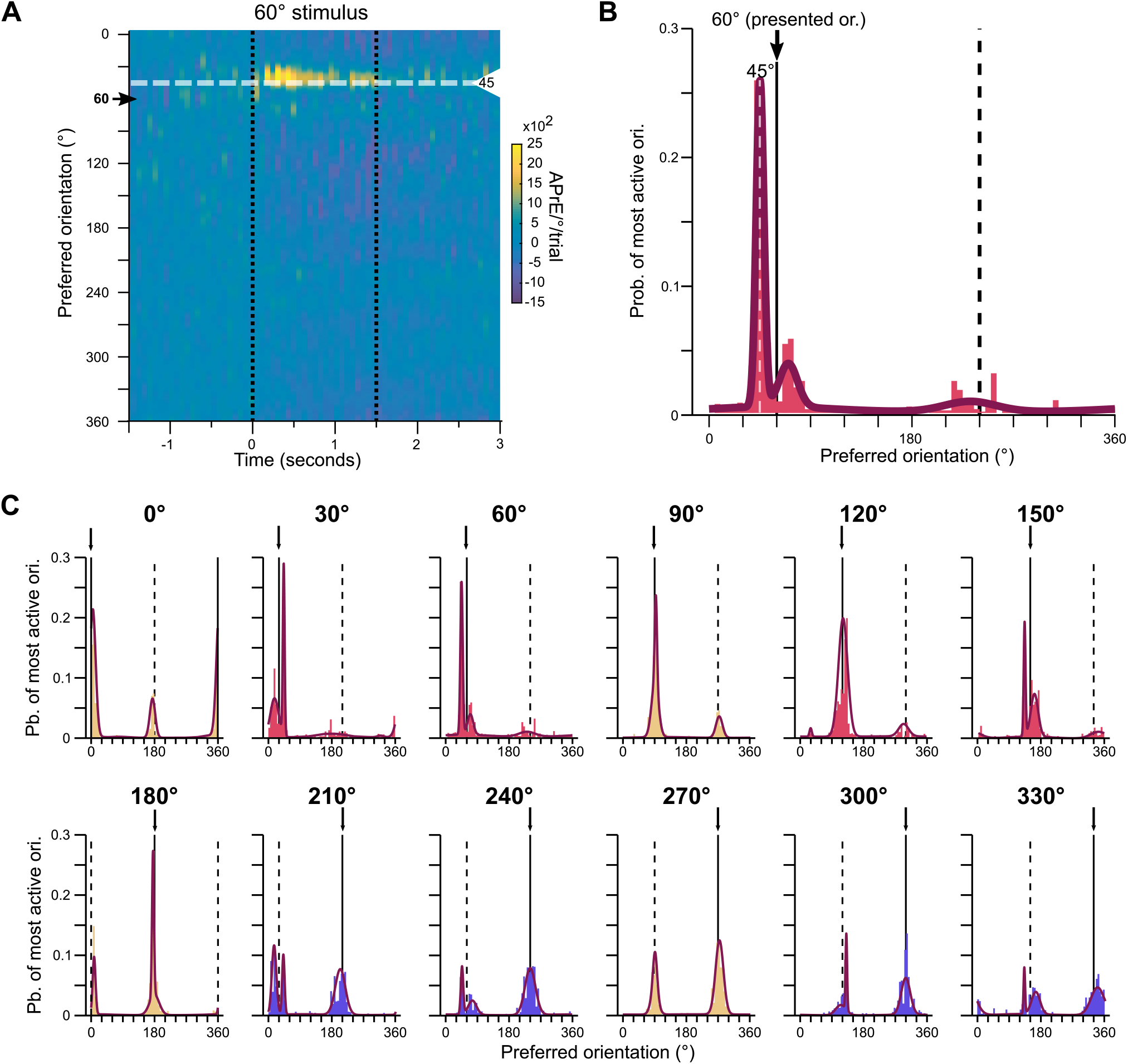
Representations of the orientations of the tuning block stimuli in the V1 of trained mice. **A**. Orientation space activity map for presentation of the 60°stimulus during the tuning block in trained mice. The arrow indicates the presented stimulus, the white dashed line indicates the orientation of the test stimulus 45°. **B**. Probability distribution of the location in the orientation space of the peak of activity evoked by the presentation of a 60°tuning block stimulus. Purple curve: three component von Mises mixture model fit. **C**. Same representation as in (B) for the 12 orientations of the tuning curve block.

**Fig. 7.**
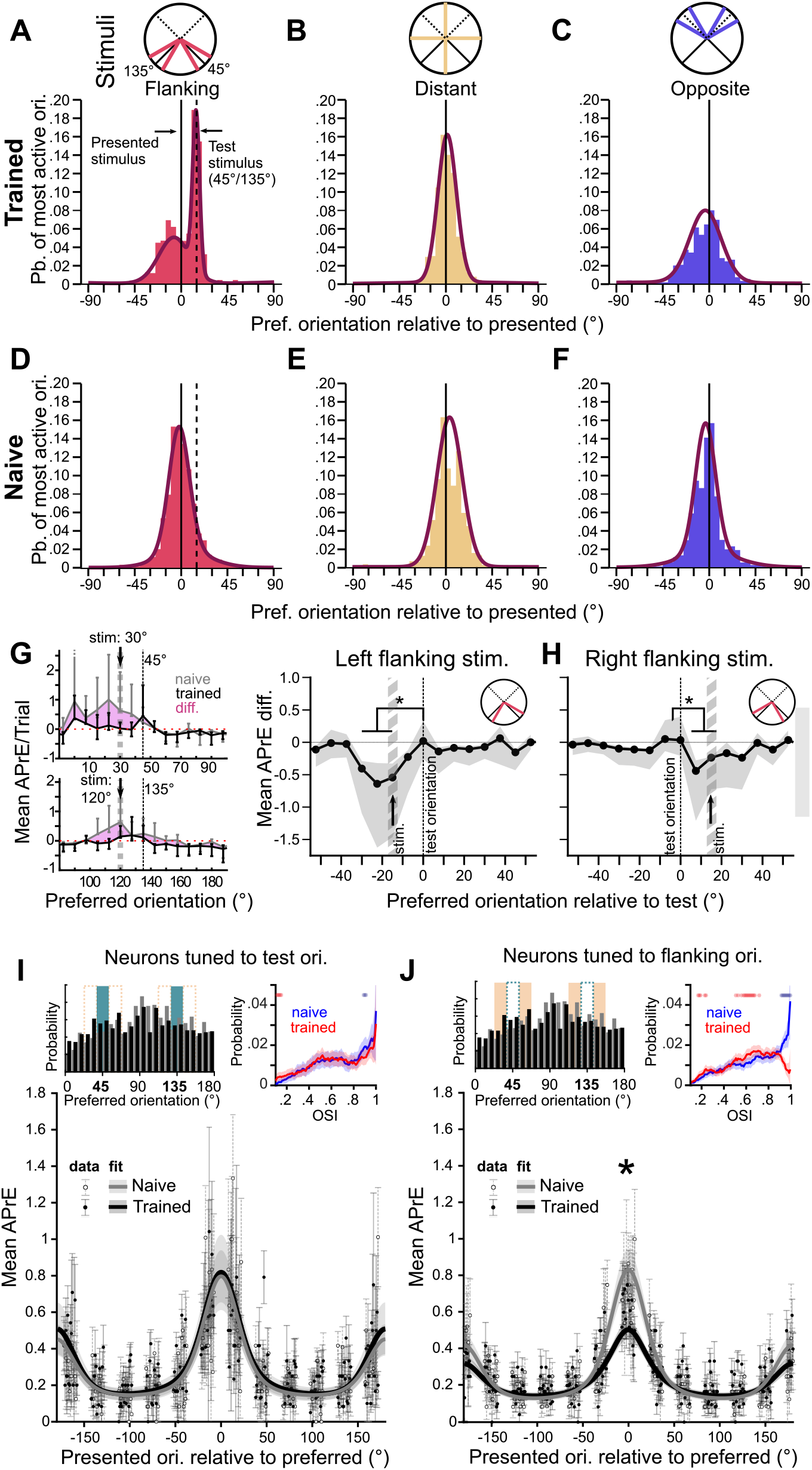
Distortion of the orientation representational space in trained mice. **A** Combination of the histograms obtained for the stimuli of the tuning curve block flanking the orientation of the test stimuli (i.e., 30°, 60° 120°, and 150°). 60°and 150°histograms were mirrored and centered such that the difference in orientation between the stimulus orientation and the test orientation was +15°. Purple curve: three component von Mises mixture model fit. **B**. Same representation as in (A) for the orientations of the tuning block distant from the orientation of the test stimuli (i.e., 0°, 90° 180°, and 270°). **C**. Same representation as in (B) for the orientations of the tuning block opposite to the orientation of the test stimuli (i.e., 210°, 240° 300°, and 330°). **D-F**. Same as A-C for naïve mice. **G**. Left: mean evoked activity for neurons binned by preferred orientation (bin size: 7.5°) for naïve (grey) and trained (black) mice. Bars indicate the s.e.m. across neurons. Shaded area shows the difference between the two profiles. Right: Difference in the neuronal responses of trained and naïve mice to the left-flanking orientations of the test stimuli (30°and 120°) as a function of preferred orientation. Shaded area indicates the 95% confidence interval obtained by bootstrapping (1,000 repetitions) **H**. Same representation as in (G) for the right-flanking orientations, i.e. 60° and 150°. **I**. Average of the tuning curves of all the neurons tuned for the test stimuli (45° and 135° ± 7.5°) of naïve (gray) and trained (black) mice. Shaded area indicates the 95% confidence interval of the mean as 1.96 s.e.m. The curves were circularly shifted such as the peak of the tuning curve would correspond to 0. Data-points represent the mean and s.e.m. of the response of individual neurons shifted along with their tuning curve fit. Top left inset: Preferred orientation distribution of the recorded neurons with the selected neurons highlighted in blue. Top right inset: distribution of Orientation Selectivity Indexes (OSI) for the neurons tuned to the test stimuli in naïve (blue) and trained (red) animals. The shaded areas indicate the 95% confidence interval obtained by bootstrapping. Stars indicate OSI values where the observed probabilities for the two groups (naïve and trained) are outside of each other’s confidence intervals. **J**. Same representation as in (I) for neurons tuned to the orientations flanking the test orientations (i.e., 30°, 60° 120°, and 150°). The star indicates a p value <0.001 for a permutation test comparing the values at the peaks of the fits between naïve and trained mice, and a small effect size (Hedges’ g >0.2).

Therefore, those neurons remained tuned to their preferred orientation as their maximum response still corresponded to that orientation, but this response was dampened compared to the response of the task neurons (which still preferred the task orientation). Altogether our results demonstrated that in trained animals, a specific suppression of flank neurons (i.e. preferring orientations flanking the orientation of the test cues) induced not only a sharpening of the test stimuli orientation representation at the population level, but also a distortion of the orientation space around the test orientations.

### Behavioral dependence of the distortion of the orientation space in trained mice

The fact that the distortion of the orientation space was only found in mice performing the task suggested that the representation changes could be specific of the behavioral expression of the learned association. To test this hypothesis, we split the trained sessions into two groups: a low D’ group made of the lower half of the D’ values (mean D’ ± s.e.m.: 1.4±0.2), and a high D’ group containing the top half performances (2.9 ± 0.1, **Fig. 8A**). For the low performance sessions, the distribution of the resampled trial activity peaks was bimodal, with maxima at −12.1 ± 1.4° and +14.7 ± 0.9° away from the orientation of the tuning block stimuli flanking the test stimuli (**Fig. 8B**). In the high D’ sessions, the peak activity of most trials was located +15.5 ± 0.6° away from the flanking stimulus orientations i.e., at the orientation of the test stimuli (**Fig. 8C**). Therefore, the likelihood that single trial representations resembled the representation of the test stimuli in trained mice was greater during high performance sessions. To better understand the link existing between distortion, learning, and performance, we examined the presence of orientation representation distortions in trained mice behaviorally disengaged from the task. We performed recording sessions in trained mice satiated for water (passive group, 7 sessions in 4 animals), 2 to 6 days (mean ± sem: 4.4 ± 1.2 days) after the last task performance (**Fig. 8D**). We found that the width of the distribution of the peak of activity evoked by the two test stimuli was comparable to that found in naïve mice (σ45°= 14.2°, C.I._95%_: [13.5 – 14.9]; σ135°= 16.9°, C.I._95%_: [16.0 – 17.7]; **Fig. 8E**, compare with Fig. 4E) and significantly broader than the distribution found in trained mice actively performing the task (compare to Fig. 4F). We then tested if the distortion of the orientation space was present in passive mice during the tuning block. We found that the larger component of the mixed model of von Mises distributions fitting the distribution of the peak of activity across trials was centered around the orientation of the displayed stimulus for orientations flanking (peak: +3.7 ± 2.4°; σ = 13.1°, C.I._95%_ : [12.5 – 13.8]; **Fig. 8F**), distant (peak: + 1.3 ± 1.2°; σ = 6.7°, C.I._95%_ : [6.3 – 7.0]; **Fig. 8G**) and opposite (peak:+4.4 ± 2.3°; σ = 12.7°, C.I._95%_ : [12.1 – 13.3]; **Fig. 8H**) to the test block stimulus orientations.

**Fig. 8.**
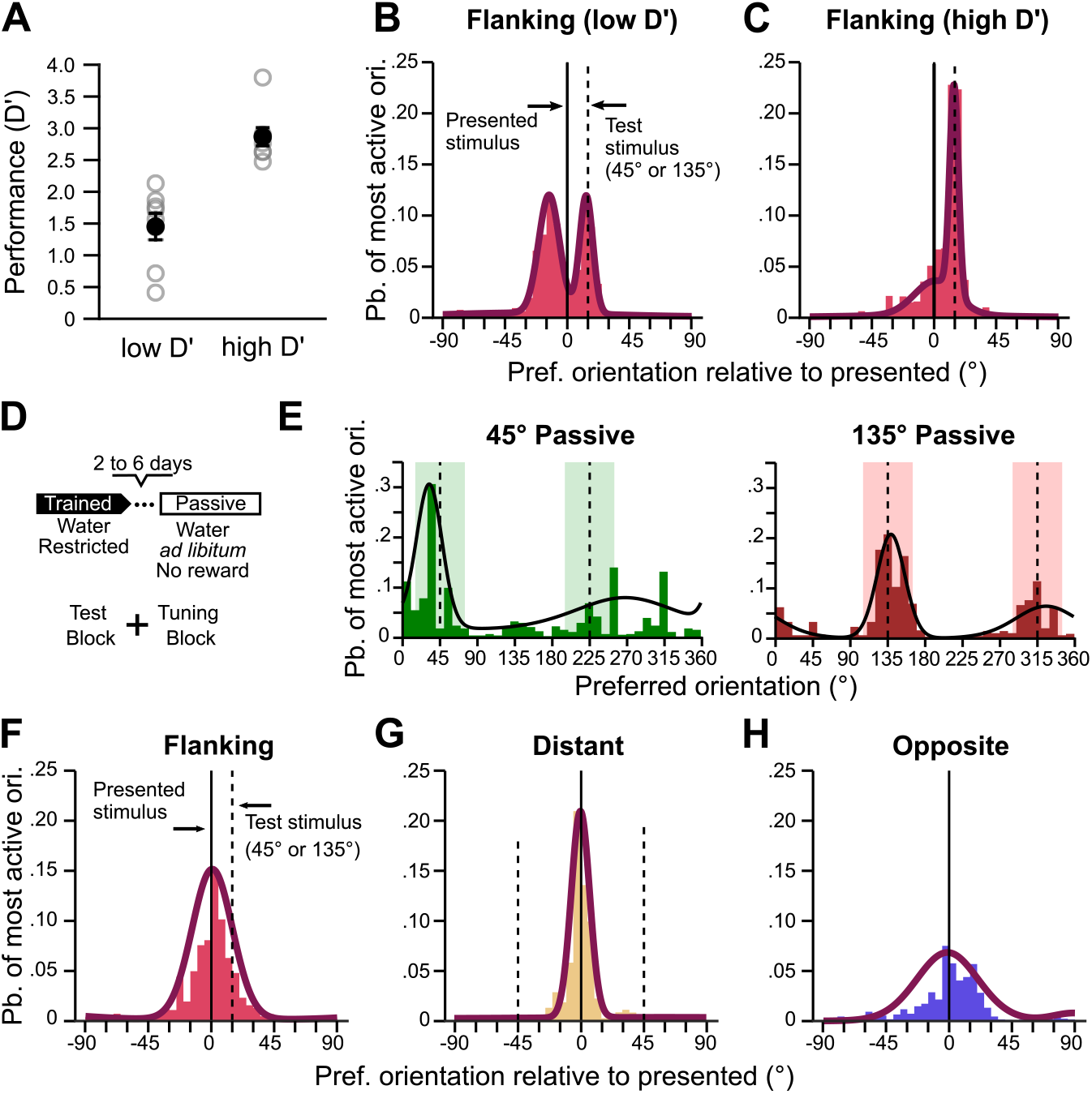
Impact of the behavioral performance on the deformation of the orientation space of trained mice. **A**. Individual D’ of sessions sorted in low or high D’ group. Black markers indicate mean and s.e.m. **B**. Same representation as in Fig. 7A but using only session of the low D’ group. **C**. Same representation as in Fig. 7A but using only session of the high D’ group. **D**. Schematic representation of the Passive group recording session. **E**. Left: Probability distribution of resampled trial activity peak in the orientation space when the 45° test stimulus was presented to passive mice. The distribution was fitted with a two component von Mises mixture model. Right: Same representation as in (A) for the presentation of the 135°test stimulus. **F**. Combination of the histograms obtained for the stimuli of the tuning curve block flanking the orientation of the test stimuli (i.e., 30°, 60° 120°, and 150°). 60°and 150°histograms were mirrored and centered such as the difference in orientation between the stimulus orientation and the test orientation was +15°. Purple curve: three component von Mises mixture model fit. **G**. Same representation as in (F) for the orientations of the tuning block distant from the orientations of the test stimuli (i.e., 0°, 90°180°, and 270°). **H**. Same representation as in (F) for the orientations of the tuning block opposite to the orientation of the test stimuli (i.e., 210°, 240°300°, and 330°).

### Saliency of the V1 responses before and after orientation discrimination training

Learning was shown to enhance the neuronal response saliency either through an increase in the proportion and/or response amplitude of neurons responsive/tuned to the task stimuli (30, 53). To determine if the distortion of the orientation space in trained mice was paired with an increase of the response saliency, we first quantified the proportions of neurons responsive to the test stimuli regardless of their orientation tuning. The fraction of neurons responsive to the 45° grating (8.1% or 149/1830) in naïve mice and in the trained mice (8.1% or 174/2141; permutation test p=0.47, Fig. 9A) were similar. A lower proportion of neurons were responsive to the unrewarded stimulus (135° drifting grating) in trained mice (4.2% or n=90/2141) than in naïve mice (8.6% or 157/1830, p<0.001; **Fig. 9A**). A similar proportion of neurons responded to both stimuli in naïve and trained mice (naïve: 3.2%, n=58/1830; trained: 2.4%, n=51/2141, p=0.053). Overall, the proportion of responsive neurons was lower in trained than in naïve mice (naïve: 16.7%, n=306/1830; trained: 12.3%, n=264/2141; p<0.001). Notably, among the responsive neurons, the relative proportion of neurons responsive to the Go and NoGo stimuli changed in favor of the Go in trained animals. We then compared the evoked activity of the responsive neurons in naïve and trained groups. For the 45° stimulus, the amount of APrEs accumulated during the first second of the stimulus presentation was slightly lower in trained mice (median APrEs/trial and 1st and 3rd quartiles, naïve: 0.81, [0.30, 1.51]; trained: 0.55, [0.22, 1.06], permutation test p=0.0015, g= 0.2, **Fig. 9B**). For the 135° stimulus, there was no significant difference between the evoked activity of the responsive neurons of the two groups (naïve: 0.64, [0.36, 1.29]; trained: 0.54, [0.21, 1.0] p=0.048, g=0.09, **Fig. 9C**). Altogether, we did not find any enhancement of the activity evoked by the task stimuli after the training, but a decrease of the population average response through a decrease of the evoked activity for the Go stimulus and of the number of responsive neurons for the NoGo.

**Fig. 9.**
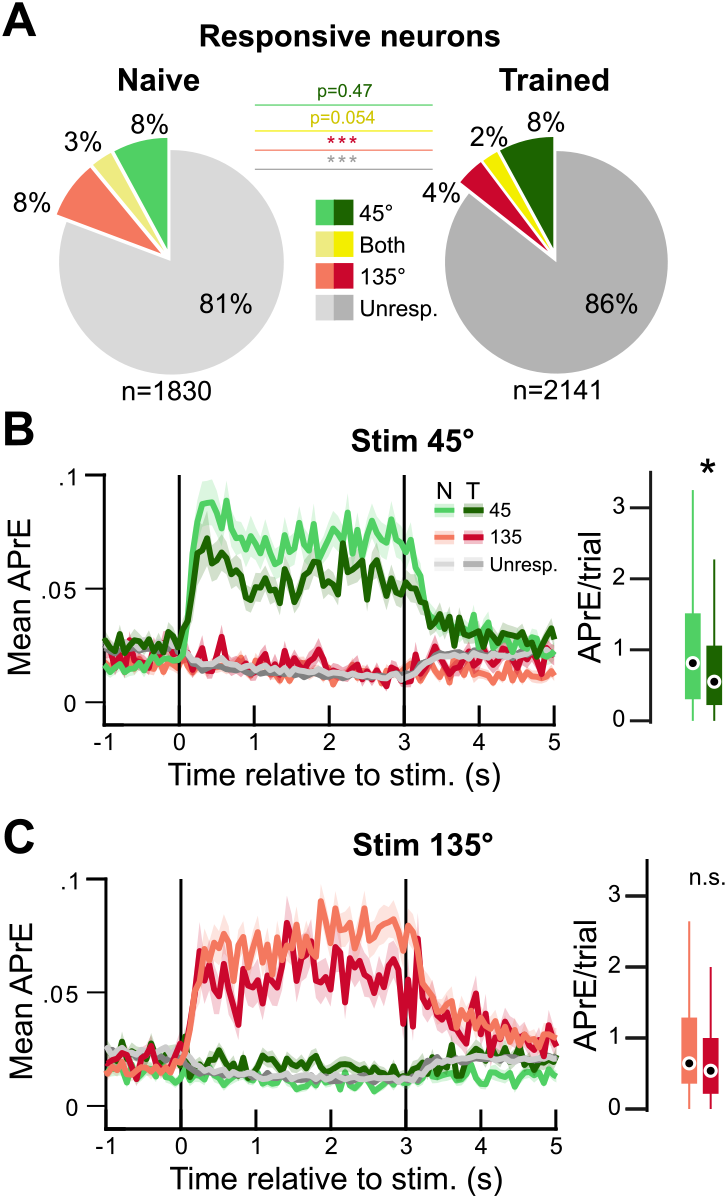
Saliency of the visually evoked responses of V1 neurons before and after training. **A**. Proportion of neurons responsive to the 45° and 135° gratings (binomial test, response vs. baseline, p<0.01). Stars indicate permutation test p-values, *** p<0.001 **B**. Time course of the response evoked by the 45°grating for neurons grouped by stimulus responsiveness. Mean across neurons ± s.e.m. Left: distribution of the sum of APrE during the first second of stimulus presentation. Box plots represent quartiles, dots indicate medians. The star indicates a significant permutation test p-value (p<0.001) and a small effect size (Hedge’s g > 0.2). **C**. Same representation as (B) but for the 135°grating presentation

## Discussion

In this study, we compared the population responses of layer 2/3 V1 neurons before and after orientation discrimination training to determine whether learning modifies the populational representation of orientations in V1. We found that: [1] the representation in V1 of the orientation of drifting gratings associated with the task was more accurate and more stable in trained mice than in naïve mice. [2] The representation of stimuli having an orientation flanking the orientations of the test cues was distorted. As a result, these oriented stimuli were represented in V1 as the test cues themselves. [3] The deformation of the orientation space in trained mice resulted from the specific suppression of the responses of neurons whose preferred orientations flanked the orientations of the task cues (**Fig. 10A,B**). [4] The transformations of the representation of orientated stimuli in V1 were context-dependent, as they were absent in mice trained to perform the task but passively viewing the cues. [5] In our behavioral paradigm, learning was not paired with an increase of the neuronal population responses to task-relevant cues.

**Fig. 10.**
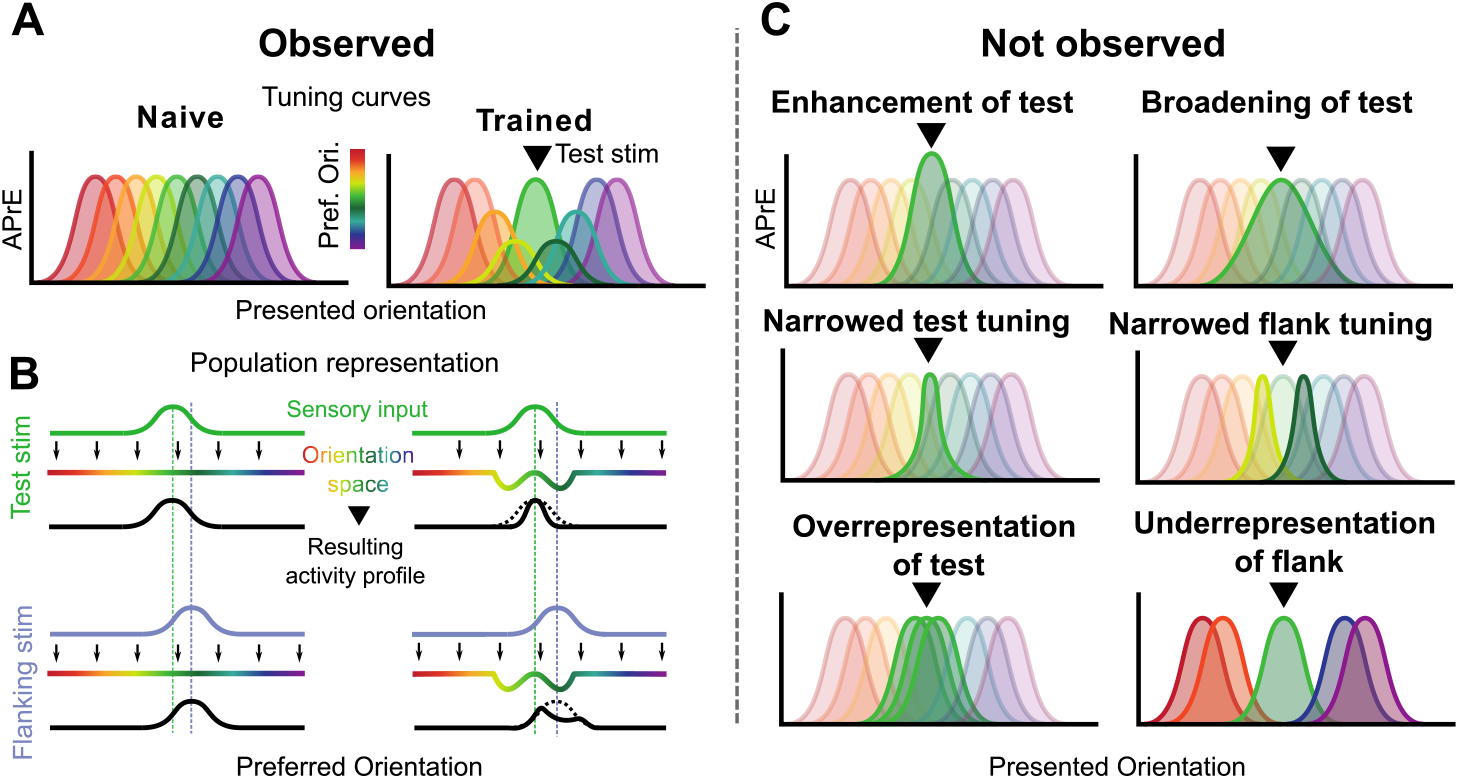
Schematic representation of the observed orientation space distortion and unobserved alternatives. **A**. Summary of the observed results. Top: tuning curves of neurons with preferences spanning the orientation space in different colors. In naïve mice (left), they are similar across orientations. In trained mice (right), neurons preferring orientations flanking that of the test have a dampened tuning curve (see Fig. 7J). **B**. Impact of these changes on orientation population representation. That schematic representation describes the population activity profile as an interaction between the sensory input and the state of the orientation space. In naïve mice (left), the amplitude of the tuning curves across preferred orientations is similar, creating a flat orientation space. When a sensory input corresponding to a given orientation is presented, the activity profile corresponds to that sensory input, for the test stimuli (top) and for the flanking stimuli (bottom). In trained mice (right), the suppression of the flank neurons creates a depression in the orientation space. When the test orientation is presented, the resulting profile is sharpened (see Fig. 3C), and when the flanking orientations are presented, the population response peaks at the location of the test orientation (see Fig. 6A-B). **C**. Potential effects of training that we did not observe in our experimental paradigm: no response enhancement of the neurons tuned to the test orientations (see Fig. 7I, Fig. 9B-C), no broadening or narrowing of their tuning curves nor that of flank neurons (see Fig. 5G, Fig. 7I-J), and no over representation (number of tuned neurons) of the test orientations or underrepresentation of the flanking orientations (see Fig. 1J, Fig. 9A).

Our main finding is that the representation of oriented stimuli in the V1 orientation space is more accurate and more reliable from trial to trial after learning. This improvement did not result from a signal increase (27) but from a reduction of the noise in the orientation space (20, 25). Indeed, the sharpening of the representation was due to the suppression of neurons tuned for flanking orientations. This suppression was still present during the tuning block presented at the end of the recording session. It provided a competitive advantage for the neurons preferring the task orientations when neighboring orientations were presented. Indeed, neurons tuned to those orientations responded less for the presentation of their own preferred stimulus than the neurons tuned for the task orientations. This suppression in the orientation space is reminiscent of the spatial modulation encountered in V1 after learning in the context of a figure-ground detection task (69) or of spatial attention (9, 10, 15, 22, 48, 49, 60, 62). Additionally, this suppression funneled the population activity toward two locations of the orientation space: the orientations of the Go and No-Go cues. It is therefore possible that this mechanism supports the capability of the visual system to generalize (2), allowing relatively small variations of the inputs to be still treated as the targets in the context of the task. If it is the case, the role of the visual cortex in the task context would be therefore to determine whether the stimulus presented is in the Go or No-Go category rather than to determine with precision the orientation of the visual stimulus. A similar phenomenon was observed in the primary auditory cortex (68), where the population activity was dynamically remodeled to represent the category of the stimuli.

The suppression of flanking neurons observed in trained mice suggests that the cellular mechanisms underpinning the changes in orientation representation found in trained mice depends on interneuron-mediated inhibition. Inhibitory interneurons have been shown to facilitate the plasticity of cortical circuits during learning (12, 19, 37, 40, 42, 55). More particularly, somatostatin neurons were shown play an important role in gating response plasticity in V1 (38). Those neurons are, at least in part, under the control of top-down signal arising from the orbitofrontal cortex (41), cingulate cortex (71) and/or retrosplenial cortex (42). Yet, parvalbumin-expressing interneurons were also shown to modulate their tuning properties after training (38), and they are associated with divisive inhibition (21) fitting what we observed (**Fig. 7J**). Furthermore, several mechanisms involving mesencephalic dopaminergic neuromodulation (4, 49, 54), and cholinergic neuromodulation (24) were also associated to perceptual learning in the context of attention and reward. Hence, it was recently proposed that cholinergic signals promote cortical plasticity during associative learning via the activation of vasoactive intestinal polypeptide-expressing (VIP) interneurons (23). Here, the changes in orientation representation were context-dependent, suggesting a pivotal role of top-down modulation. However, we cannot exclude the participation of local synaptic plasticity (53). Therefore, the respective role of top-down modulation, local synaptic plasticity, GABAergic inhibition, and neuromodulation in the distortion of the orientation space described in this study will need to be further investigated in the future.

Contrary to previous reports that showed a specific enhancement of the rewarded cue representation (28, 30, 59, 61), we observed similar changes of the orientation representation of the rewarded and non-rewarded cues. Moreover, we did not find any upregulation of the responses supporting an enhancement of to the task stimuli representation (30, 36, 53) nor individual tuning width modulation ((28), **Fig. 10C**). Discrepancies between studies investigating the impact of learning on sensory processing are common. This variability is likely due to the existence of differences between the experimental design of those studies (67). The type of task (operant chamber (28), classical conditioning (36), or Go/NoGo task in a virtual reality set up where the stimulus presentation is coupled with locomotion (30, 53) and the nature of the reward and/or the punishment likely affect the perceived value of the task stimuli, changing the nature and magnitude of reward-related signaling in the brain (57). Opposite effects of learning on stimuli representation in A1 for the same task can be induced by forcing the animals to use alternative strategies (8). Moreover, visual stimuli features can affect the susceptibility of V1 neurons to present task-related modulations. For instance, the size of the stimulus is known to influence neuronal responses in V1: full-field drifting gratings trigger surround inhibition that enhances the sparseness of the V1 population response compared to more localized stimuli (1, 66). The contrast of visual stimuli also affects context-dependent modulations (33, 46). In this study, we chose a task that mice acquire easily, and there should be a priori no need to improve the representations of the stimuli in V1 to perform it. The angular distance between the Go and NoGo cues is maximal (90°), we use high-contrast (75%) full-field stimuli, and mice are not required to navigate in a virtual reality environment. Hence, the representation of the 45° and 135° stimuli are already highly distinct in naïve mice V1, as they do not overlap in the orientation space (Fig. 3A-D). This suggest that the representations of the two stimuli were sufficiently separable for naïve mice to discriminate them reliably. Therefore, the computational challenge of our task is likely to be more centered on the association of the sensory inputs with the appropriate responses, rather than on a perceptual improvement. The structure of the changes we observed, i.e., sparser and more consistent population responses both for Go and NoGo, can fit those computational requirements by promoting associative learning mechanisms such as Hebbian plasticity (51). Finally, it is possible that the presence of audiovisual blocks (excluded from the analysis) influenced the results as the presence of sound was shown to improve the representation of oriented stimuli in naïve mice (45). However, the improved representation of the test stimuli and the deformation of the orientation space evidenced in trained mice cannot be attributed to the mere presence of sound as audiovisual blocks were also present in the session of naïve and passive mice. Further experiments will be necessary to determine if the presence of audiovisual cues during the recording sessions could have had some synergic effect with the phenomenon described here.

To conclude, our results provide new evidence that, during the performance of an orientation discrimination task, visual processing is biased toward the cues of the task. This bias effectively structures the neuronal responses around attractors in the encoding space. It is possible that this mechanism facilitates learning by allowing the generalization of the properties of the cue of interest to a larger proportion of the encoding space. Because we cannot find those attractors in passive mice, our results also suggest that the deformation of orientation representational space that results from learning is an ad-hoc modulation. The function of this filter imposed transiently to the sensory cortices (possibly by top-down signals), would be to facilitate the extraction of task-relevant information.

## AUTHOR CONTRIBUTIONS

J.M. performed the experiments; J.C. designed and performed the analysis with contributions from P-O.P. and B.E.; P-O.P. designed and supervised the study. J.C. and P-O.P. wrote the manuscript with feedbacks from all the co-authors.

## ACKNOWLEDGEMENTS

This work was funded by the Whitehall Foundation (grant 2015-08-69), the Charles and Johanna Busch Biomedical Grant Program, and the National Institutes of Health – National Eye Institute (Grant #R01 EY030860). J.C. was supported by a Fyssen Foundation postdoctoral fellowship. The authors are grateful to Denis Paré, Bart Krekelberg, Drew Headley and the members of the laboratory for their comments on the manuscript

